# Molecular basis for SARS-CoV-2 spike affinity for human ACE2 receptor

**DOI:** 10.1101/2020.09.10.291757

**Authors:** Julián M. Delgado, Nalvi Duro, David M. Rogers, Alexandre Tkatchenko, Sagar A. Pandit, Sameer Varma

## Abstract

Severe acute respiratory syndrome coronavirus-2 (SARS-CoV-2) has caused substantially more infections, deaths, and economic disruptions than the 2002-2003 SARS-CoV. The key to understanding SARS-CoV-2’s higher infectivity may lie in its host receptor recognition mechanism. This is because experiments show that the human ACE2 protein, which serves as the primary receptor for both CoVs, binds to CoV-2’s spike protein 5-20 fold stronger than SARS-CoV’s spike protein. The molecular basis for this difference in binding affinity, however, remains unexplained and, in fact, a comparison of X-ray structures leads to an opposite proposition. To gain insight, we use all-atom molecular dynamics simulations. Free energy calculations indicate that CoV-2’s higher affinity is due primarily to differences in specific spike residues that are local to the spike-ACE2 interface, although there are allosteric effects in binding. Comparative analysis of equilibrium simulations reveals that while both CoV and CoV-2 spike-ACE2 complexes have similar interfacial topologies, CoV-2’s spike protein engages in greater numbers, combinatorics and probabilities of hydrogen bonds and salt bridges with ACE2. We attribute CoV-2’s higher affinity to these differences in polar contacts, and these findings also highlight the importance of thermal structural fluctuations in spike-ACE2 complexation. We anticipate that these findings will also inform the design of spike-ACE2 peptide blockers that, like in the cases of HIV and Influenza, can serve as antivirals.

## Introduction

Within ten months of its emergence, the severe acute respiratory syndrome coronavirus-2 (SARS-CoV-2) has caused more than 23 million confirmed infections and over 800,000 deaths globally, and these infections continue to grow rapidly [1]. In contrast, its genetic variant, SARS-CoV, which caused the 2002-2003 outbreak, led to far fewer infections and it was relatively easier to contain, although it presumably had a much higher fatality rate [2]. The underlying reason for CoV-2’s relatively higher infectivity, however, remains unknown [2].

The key to understanding CoV-2’s higher infectivity in humans may lie in its host receptor recognition mechanism. This is because of the following. Both CoV and CoV-2 use the angiotensin converting enzyme 2 (ACE2) as their primary modes of attachment and entry into human cells [3]. They bind to human ACE2 receptors using their respective transmembrane spike proteins. However, surface plasmon resonance and bilayer interferometry experiments show that ACE2 binds to CoV-2’s spike protein 5-20 fold stronger than CoV’s spike protein (Table S1 of supporting information) [4–7], which may, at least partly, explain its higher infectivity.

The molecular basis underlying the different binding affinities of CoV and CoV-2 spike proteins to ACE2, however, remains unknown. In fact, comparison of high-resolution X-ray structures of their spike-ACE2 complexes [4, 6, 8] leads to a different proposition. The receptor binding domains (RBDs) of the spike proteins of CoV and CoV-2 have almost similar structures, and they interact with almost identical regions of the proteolytic domain (PD) of ACE2. Out of the forty three amino acid that are different in the spike RBDs of CoV and CoV-2 strains used for structure determination, ten are at the ACE2 binding region [4]. These ten amino acid differences do not alter the numbers of hydrogen bonds and salt bridges at the spike-ACE2 interface [4]. They only make CoV’s spike protein to have a more extensive hydrophobic contact with ACE2, which should have, in principle, made CoV’s spike protein bind more strongly than CoV-2’s spike protein, rather than the opposite [9].

Recent studies show that thermal fluctuations in structure can affect virus-host interactions and viral entry [10], and perhaps they may also explain why ACE2 binds more tightly to CoV-2’s spike protein. To examine this and go beyond insights obtained from X-ray structures, here we carry out a comparative analysis of spike-ACE2 complexes using all-atom molecular dynamics (MD) simulations at physiological temperature and in explicit solvent. Additionally, we carry out free energy calculations to examine the effect of mutating selected spike residues on spike-ACE2 binding affinity. Results from these studies provide an atomically-detailed basis for why ACE2 binds to CoV-2 spike more strongly compared to CoV spike. This structural and energetic data will be useful to groups designing small molecules, polymers, and antibodies targeting the spike-ACE2 interaction.

## Results and Discussion

From 3 μs long MD trajectories of ACE2 PD complexed with spike RBDs of CoV and CoV-2, we first examine differences in binding modes, polar contacts, hydrophobic contacts and interfacial waters. Next, using conformations selected from MD simulations, we examine the effect of mutating selected CoV spike RBD residues on spike-ACE2 binding free energy.

### Comparison of MD ensembles

#### Binding modes

To characterize spike-ACE2 binding modes, we extract from each simulation spike-ACE2 conformations at one nanosecond intervals. This yields 3001 conformations for each complex. We calculate root mean square deviations (RMSDs) between each of the (3001 × 3000)/2 conformation pairs. For calculating RMSDs, we consider the backbone atoms of only those amino acids in ACE2 and spike that are part of the spike-ACE2 interface. The interface is defined geometrically, and an amino acid is considered to part of the interface if any of its heavy atoms is within 5 Å from the complementary protein in any of the 3001 conformations. These pairwise RMSDs are shown in Figure 1. These RMSDs are then taken as a measure of similarity in the affinity propagation algorithm [11, 12], which clusters these conformations into 5 and 6 groups, respectively, in the CoV and CoV-2 spike-ACE2 systems. The advantage of the affinity propagation algorithm over traditional clustering approaches is that it does not assume *a priori* the number of clusters or a cutoff value for delineating clusters. We adopted this unsupervised machine learning algorithm previously to cluster correlations in structural fluctuations [13].

**Figure 1.**
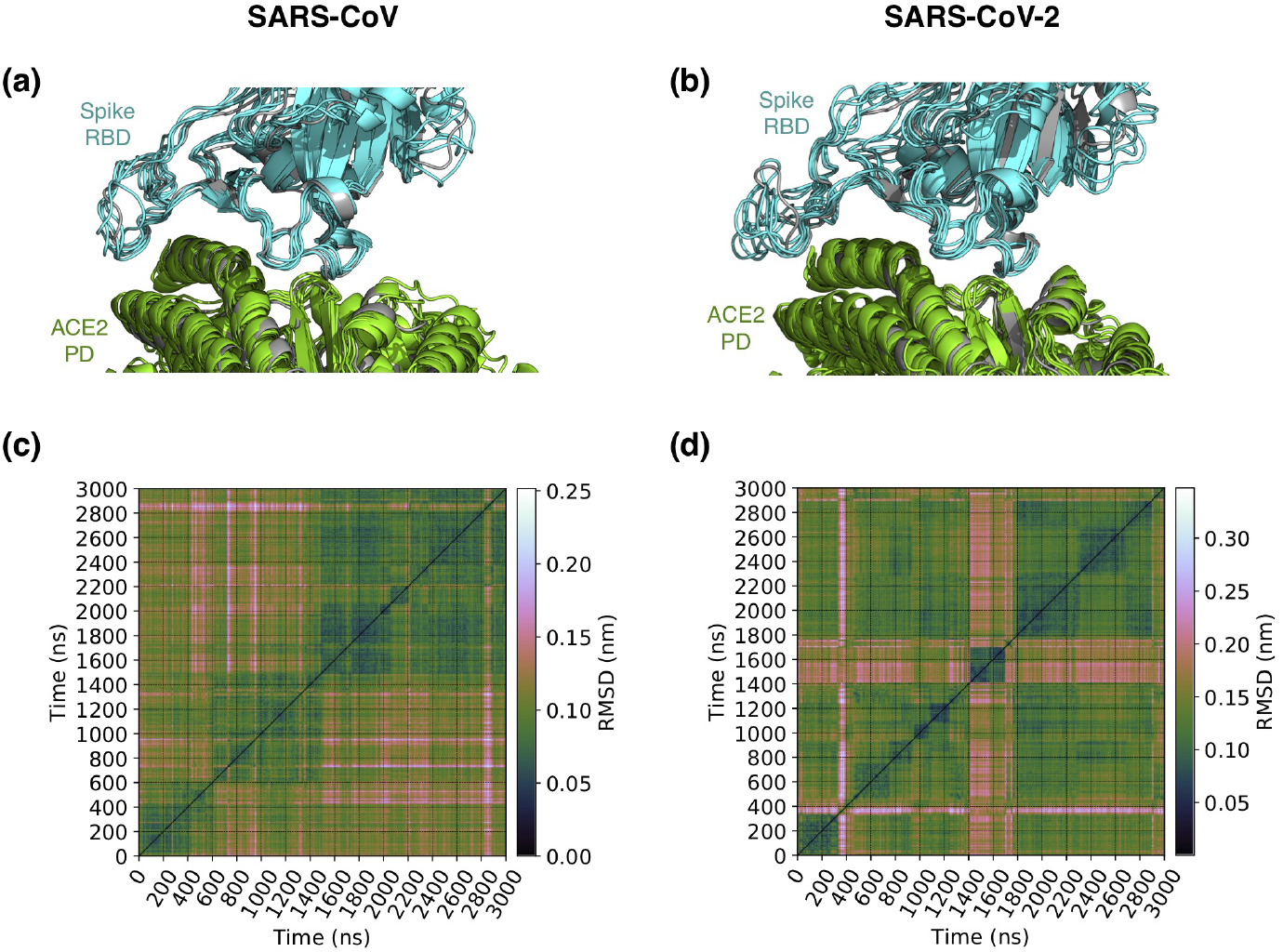
Binding modes of spike-ACE2 complexes in MD simulations. **(a)** and **(b)** show, respectively, the distinct binding mode conformations of the complexes containing spike proteins of CoV and CoV-2. These conformations are superimposed over the X-ray structures of spike-ACE2 complexes (grey). They are the lowest energy conformations of the five and six binding modes identified, respectively, for the complexes involving CoV and CoV-2 spike proteins. Binding modes are identified by clustering conformations extracted every nanosecond from MD. Conformational clustering is performed using affinity propagation [11,13] in which we take RMSD as an index of similarity between conformations. **(c)** and **(d)** show these pairwise RMSDs.

The lowest energy conformation of each of the 5 and 6 clusters of CoV and CoV-2 spike-ACE2 complexes are shown in Figure 1. We note that the overall topologies of these cluster representatives, or binding modes, closely resemble their respective X-ray structures. The main variation in the binding modes is in the structures of the spike RBD loops at the binding interface.

#### Local spike-ACE2 contacts

To examine differences in local interactions, we first determine hydrogen bonds and salt bridges between spike RBD and ACE2 PD. Hydrogen bonds are computed using the geometric definition proposed by Luzar and Chandler [14]. Salt bridges are defined using a 4 Å cutoff between carboxyl carbon and amine/guanidine nitrogen. The choice of this cutoff distance is discussed in Figure S1 of the supporting information. Figure 2a compares time evolutions of inter-protein hydrogen bonds and salt bridges. The key observation we make is that, on average, ACE2 engages in discernibly greater numbers of hydrogen bonds and salt bridges with CoV-2’s spike RBD compared to CoV’s spike RBD. This difference was not apparent from comparison of X-ray structures, where both complexes were reported to have equal numbers of salt bridges and hydrogen bonds [4].

**Figure 2.**
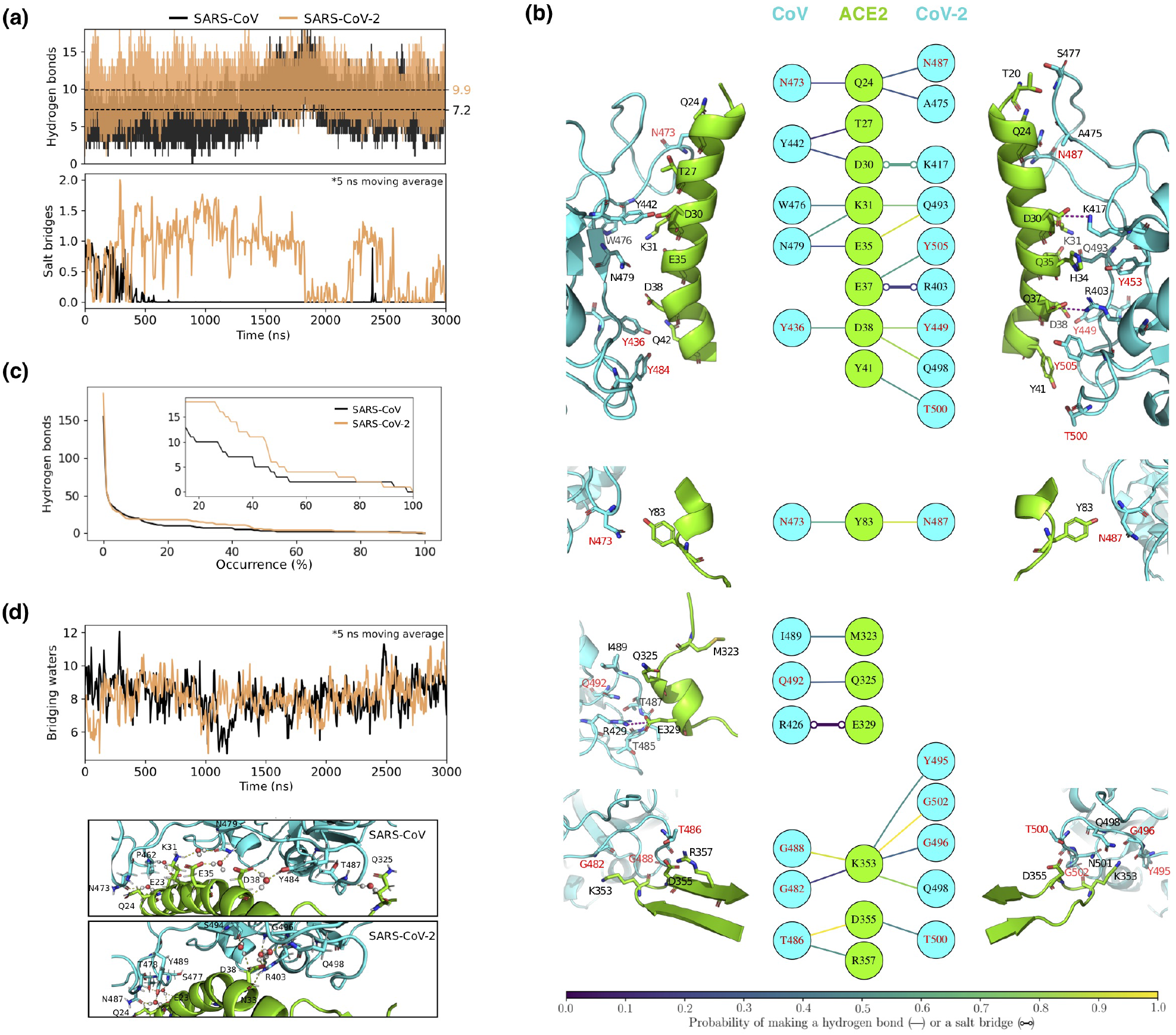
Polar contacts of ACE2 with spike proteins of CoV and CoV-2. **(a)** Time evolution of hydrogen bonds and salt bridges between ACE2 and spike proteins. Dashed lines indicate time-averages. **(b)** Structural map of hydrogen bonds and salt bridges between ACE2 and spike. ACE2 associates with spike at four regions that are non-contiguous in its primary sequence. These four interfacial regions are shown separately. The colors of the lines connecting the residues in the central panel indicate their occurrence probabilities. Note that for the sake of clarity, only those hydrogen bonds and salt bridges are shown that are observed for at least 15% of the total simulated time. The amino acid of spike labeled in red are the ones that are conserved in the two spike RBDs. **(c)** Numbers of unique hydrogen bonds as functions of their occurrence probabilities. The inset zooms in on the 15-100% probability region. **(d)** Time evolution of waters that bridge interactions between ACE2 and spike by hydrogen bonding simultaneously with both proteins. The image below show the time evolution plot shows the bridging waters in the 3 *μ*s snapshots of the MD trajectories.

To gain further insight, Figure 2b compares the structural maps of the spike-ACE2 hydrogen bonds and salt bridges. Several of the polar contacts observed in MD are also noted in X-ray structures [4, 6, 8], and, in fact, we note that MD simply yields a higher combinatorics of contacts than those discerned from X-ray structures (Fig. S2 of supporting information). We attribute this difference between MD and X-ray structures to structural thermal fluctuations present in MD. This difference may also be attributed to the dynamics of interfacial water molecules [15], which is present in MD, but absent in X-ray structures.

From Figure 2b, we also note that the higher numbers of polar interactions in CoV-2’s spike-ACE2 interface do not emanate from one specific region of the interface. In fact, almost all residues of CoV-2 spike, including those that are conserved in CoV spike, have a higher probability of forming hydrogen bonds (or salt bridges) with ACE2. The probabilities here refer to the fractions of frames in which amino acids are found to be hydrogen bonded (or salt-bridged). To examine these hydrogen bond probabilities collectively, we plot in Figure 2c the numbers of unique hydrogen bonds as functions of their occurrence probabilities. We note that CoV-2’s spike-ACE2 interface has not only more possible combinations of hydrogen bonds compared to CoV-2’s spike-ACE2 interface, but the net probability of formation of hydrogen bonds is also higher in the CoV-2 spike-ACE2 interface.

Polar interactions between spike and ACE2 can also be bridged by interstitial water molecules, that is, water molecules can hydrogen bond simultaneously with both proteins. These bridging waters are considered to stabilize protein-protein interactions [16]. We note from Figure 2d that the spike-ACE2 interfaces of CoV and CoV-2 have similar numbers of bridging waters, which supports the possibility that bridging waters have comparable effects on the stabilities of both spike-ACE2 complexes.

Finally, consistent with X-ray structures of the spike-ACE2 complexes, we note that ACE2 makes a more extensive hydrophobic contact with the spike RBD of CoV (Fig. S3 of Supporting Information). A higher hydrophobic contact typically implies a higher binding affinity [9]. Therefore, this interaction should drive binding of ACE2 in favor of CoV, and not CoV-2.

### Effect of spike mutations on binding free energy

The spike RBDs of CoV and CoV-2 strains that were used for structure determination contain forty three amino acid differences. Ten of these amino acid differences are at the ACE2 binding region [4]. This raises the question of the extent to which the differences noted above in local interactions result from differences in local chemistries at spike-ACE2 interfaces. This is important to know because changes in amino acids distant from the interface can affect protein-protein binding substantially, and even in the absence of discernible structural change [17]. To address this, we substitute simultaneously eight of these amino acids in CoV spike RBD to their corresponding types in the CoV-2 spike RBD (Figure S4 of supporting information), and determine the effect of this mutation on spike-ACE2 binding free energy. We do not engineer the mutation L443→F456 because neither L443 in CoV nor its corresponding residue F456 in CoV-2 is found to make hydrophobic contact with ACE2 (Figure S3 of Supporting Information). The mutation P462→A475 is also not engineered due to the current unavailability of established transition pathways [18].

The eight mutations that we engineer in CoV spike RBD are V404K, R426N, T433G, Y442L, L472F, N479Q, T484Q and T487N. Note that these mutations leave the net charge, and also the ratio of hydrophobic and polar side chains at the interface unaltered. The expectation is that if these local chemical differences indeed explain CoV-2’s higher binding affinity for ACE2, then these mutations should increase the binding free energy of CoV spike to ACE2. Free energy calculations show that these mutations do increase the binding affinity of CoV spike RBD by 5.4 ± 0.4 kcal/mol (Figure S4 of supporting information).

Note, however, that this estimate is larger than the experimental range of 1-2 kcal/mol determined for the binding free energy difference between CoV and CoV-2 spike-ACE2 complexes [4–7]. This could be due to two reasons. Firstly, this may partly result from inaccuracy in the employed force field, although, based on recent studies [18], the extent of this error is expected to be around 1-2 kcal/mol. Secondly, this deviation may partly be due to correlated and allosteric effects that originate from the 30+ amino acids in CoV spike RBD that differ from CoV-2 spike RBD, and are not mutated in CoV spike RBD. This possibility is supported by the data in Figure 2b, where we note differences even in the ACE2 interactions of amino acids that are conserved in CoV and CoV-2 spike RBDs. For example, the hydrogen bond probability of Y83 in ACE2 with N473 in CoV spike is distinctly smaller compared to its hydrogen bonding probability with N487 in CoV-2 spike.

## Conclusions

Analysis of MD trajectories yields three main differences between the spike-ACE2 complexes of CoV and CoV-2. Firstly, consistent with observations from X-ray structures, ACE2 makes a more extensive hydrophobic contact with CoV’s spike. This should, in principle, drive binding of ACE2 in favor of CoV’s spike rather than CoV-2’s spike. Secondly, there are distinctly greater numbers of hydrogen bonds and salt bridges in CoV-2’s spike-ACE2 complex. Finally, the combinatorics as well as the individual and net probabilities of these polar contacts are higher in CoV-2’s spike-ACE2 complex. The latter two differences will drive ACE2 binding in favor of CoV-2, and also implicate thermal fluctuations in structure to be important to selective spike-ACE2 complexation. These observations lead to the conclusion that the higher affinity of CoV-2’s spike to ACE2 is due to higher numbers and probabilities of polar contacts, which also compensate for the more extensive hydrophobic contacts in CoV’s spike-ACE2 interface. The caveat here is that local interactions are key to driving specificity. In fact, free energy calculations directly support this caveat, where substitutions of eight interfacial amino acids in CoV spike to corresponding ones present in CoV-2’s spike increase its binding affinity to ACE2. Additionally, since these eight amino acids collectively have the same charge and similar hydrophobic/polar chemistry ratios, we also conclude that the specific structural locations of these amino acids matter. As such, results from both free energy calculations and equilibrium MD simulations do suggest the role of allosteric and correlated effects in spike-ACE2 complexation.

Given existing evidence on the general reliability of employed MD methods [13, 18–20], and our recent work [21] that yielded validated predictions on virus-host protein-protein interactions [22], we consider our qualitative conclusions to be robust. Nevertheless, from the perspective of intermolecular interaction theory, the underlying potential energy functions that we employ do rely on describing interactions using point charges, no polarization, and only pairwise vdW interactions. These approximations should be properly scrutinized in future studies of complex protein-protein interactions. Higher-level electronic structure calculations are in progress to assess the role of multipole electrostatics, induced polarization, and many-body dispersion interactions on spike-ACE2 binding.

Overall, the molecular understanding that this work provides on the relative binding affinities of CoV and CoV-2 spike to ACE2 is important for understanding their different infectivity rates. Additionally, it is also expected to lend direct insight into designing spike-ACE2 blockers that, like in the cases of HIV and Influenza, can serve as antivirals [23].

## Methods

### Molecular dynamics

All MD simulations are performed using Gromacs 2020 [24]. Protein and water bonds are restrained [25, 26], and consequently an integration time step of 2 fs is employed. Simulations are conducted under isobaric-isothermal boundary conditions. Pressure is regulated at 1 bar using a coupling constant of 1 ps and a compressibility of 4.5 × 10^−5^ bar^−1^. Temperature is maintained at 310 K. Extended ensemble approaches are used for maintaining both temperature and pressure [27–29]. Electrostatic interactions beyond 10 Å are computed using the particle mesh Ewald scheme [30] with a Fourier grid spacing of 1.5 Å, a fourth-order interpolation. van der Waals interactions are computed explicitly for interatomic distances smaller than 10 Å. We use Amber99sb-ILDN parameters to describe protein and ions [19], and SPC/E parameters to describe water molecules [31]. This force field has been demonstrated to perform well in reproducing structural data from X-ray diffraction and NMR spectroscopy [19, 20], and also dynamics data from NMR spectroscopy [13]. The ACE2 protein contains a Zn^2+^ ion in its catalytic core, which is about 20 Å away from the spike-ACE2 interface. In line with earlier work on modeling Zn^2+^ ions in proteins [32], the coordination of the Zn^2+^ ion in ACE2 is maintained through application of distance restraints. Specifically, flat-bottomed quadratic potentials are assigned to distances between the ion and three atoms of ACE2, H374/NE2, H378/NE2 and E402/OE1, that are observed to coordinate it in X-ray structures. In MD simulations, the average restraining energies for the CoV and the CoV-2 complexes are found to be small, that is, 0.14 ± 0.08 and 0.11 ± 0.06 kcal/mol, respectively.

Starting coordinates of CoV’s spike-ACE2 complex are taken from its X-ray structure (PDB ID: 2AJF) [8], and those of CoV-2’s spike-ACE2 complex are taken from its cryo-EM structure (PDB ID: 6M17) [33]. Note that a higher resolution X-ray structure of CoV-2’s spike-ACE2 complex is now also available [4], and is used in our analysis, but was unavailable at the start of this project. Nevertheless, there is only little difference between the X-ray and cryo-EM structures - the RMSD between all heavy atoms is < 1 Å, which is less than the RMSDs between the different interfacial binding modes observed in MD simulations. The carbohydrate groups in spike are removed, as they have been shown to have no effect on CoV’s spike binding to ACE2 [34]. The missing loops in the ACE2 protein in 2AJF are built using MODELLER [35]. To make ACE2 PD sequences identical in CoV and CoV-2 constructs, the N-terminal residues 19 and 20 in ACE2 of 6M17 are built, and the C-terminal of ACE2 in 6M17 is truncated at the last C-terminal residue resolved in 2AJF. The N- and C-termini of both spike RBD and ACE2 PD are capped with ACE and NME, respectively. Hydrogen atom positions and histidine types are determined using the PDB2PQR algorithm [36]. Each of the two complexes is initially placed in a cubic unit cell of length 160 Å, and then energy minimized using the steepest descent algorithm implemented in Gromacs. The vacant space in the box is then filled with water, and the system is again subjected to energy minimization. The unit cells containing CoV and CoV-2 spike-ACE2 complexes contain, respectively, 131,995 and 131,897 waters. Note that the crystallographically resolved waters are retained. Na^+^ and Cl^−^ ions are added by randomly substituting non-crystallographic waters. NaCl concentration is set at 50 mM with extra Na^+^ ions to compensate for the charge of the complex. Specifically, the CoV spike-ACE2 unit cell contains 144 Na^+^ and 120 Cl^−^ ions, the CoV-2 spike-ACE2 unit cell contains 143 Na^+^ and 120 Cl^−^ ions. After adding salt, the system is energy minimized a final time. Each of these two complexes is then subjected to 3 *μ*s long MD simulations.

In addition to carrying out MD simulations of spike-ACE2 complexes, we also carry out 0.5 *μ*s long MD simulations of isolated spike RBDs of CoV and CoV-2 in solution. These simulations are used for getting starting conformations for free energy calculations. For these simulations, the starting coordinates of spike RBDs are taken from final snapshots of the MD simulations of the spike-ACE2 complexes. Each of the two spike RBDs is initially placed in a cubic unit cell of length 90 Å, and the protocol described above is followed for adding waters and salt. The unit cell containing CoV’s spike RBD contains 23247 waters, 21 Na^+^ and 22 Cl^−^ ions, and the unit cell containing CoV-2’s spike RBD contains 23240 waters, 21 Na^+^ and 23 Cl^−^ ions.

### Free energy calculations

The effect of mutating spike residues on its binding free energy with ACE2 is determined as

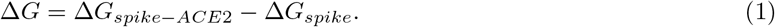

Here, Δ*G*_*spike-ace*2_ is the effect of mutations on the free energy of the spike-ACE2 complex in solution, and Δ*G_spike_* is the effect of mutations on the free energy of isolated spike in solution. Δ*G_spike_* and Δ*G*_*spike-ace*2_ are computed using thermodynamic integration, and using the 5-point Gauss-Quadrature rule, that is,

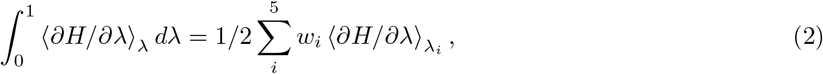

where the weights *w_i_* = {0.237,0.479,0.559,0.479,0.237} and λ_*i*_ = {0.047,0.231,0.5, 0.769, 0.953}.

The starting conformations for engineering mutations are taken from MD simulations. Hybrid topology files, which contain coordinates and force field parameters for both states of the amino acids (natural and mutated) are constructed using the PMX module [18]. To avoid singularities and numerical instabilities that may arise due to particle appearance and annihilation, we use a modified form of the “soft core” potentials suggested by Beutler et al. [37] implemented in Gromacs. In these soft core potentials, the distances between particles ‘*i*’ and ‘*j*’ in state A (λ = 0) are modified as 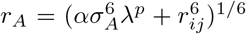 and those between particles in state B (λ =1) are modified as 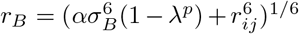. In these expressions, *σ* = (*C*_12_/*C*_6_)^1/6^ is the ratio of the LJ parameters, and if either *C*_12_ or *C*_6_ is zero, then we take *σ* = 3 Å. We set the soft core parameters to be *α* = 1 and *p* = 1. Sampling is conducted using stochastic dynamics and under NVT conditions. For each λ_*i*_, *∂H*/*∂*λ is averaged for 250 ns (Figure S4 of Supporting Information), and standard errors are determined from the final 50 ns using block averaging.

## Acknowledgments

This research used resources of the Oak Ridge Leadership Computing Facility at the Oak Ridge National Laboratory, which is supported by the Office of Science of the U.S. Department of Energy under Contract No. DE-AC05-00OR22725.

## Notes

### Competing Interest Statement

The authors have declared no competing interest.

