## Supporting Information for "Molecular basis for SARS-CoV-2 spike affinity for human ACE2 receptor"

Table S1: Comparison of ACE2-spike binding affinities from different experiments. $k_{\mathrm{on}}$, $k_{\mathrm{off}}$ and $k_d$ are, respectively, the association rate constants, dissociation rate constants and the equilibrium dissociation constants. SPR and BI are abbreviations for surface plasmon resonance and bilayer interferometry, respectively. [a]taken from reference,[1] [b]taken from reference,[2] [c]taken from reference,[3] [d]taken from reference.[4]

| Experiment | SARS-CoV | | | SARS-CoV-2 | | |
|---|---|---|---|---|---|---|
| | $k_{\mathrm{on}}$ ($\times 10^5$ 1/Ms) | $k_{\mathrm{off}}$ ($\times 10^{-3}$ 1/s) | $k_d$ (nM) | $k_{\mathrm{on}}$ ($\times 10^5$ 1/Ms) | $k_{\mathrm{off}}$ ($\times 10^{-3}$ 1/s) | $k_d$ (nM) |
| SPR[a] | 13.67 | 43.17 | 31.59 | 14.0 | 6.544 | 4.674 |
| SPR[b] | 3.62 | 112 | 325.8 | 1.88 | 2.76 | 14.7 |
| SPR[c] | 2.01 | 37.0 | 185 | 1.75 | 7.75 | 44.2 |
| BI[d] | 1.4 | 0.71 | $5.0 \pm 0.1$ | 1.4 | 0.16 | $1.2 \pm 0.1$ |

Figure S1: **Geometric definition of salt bridge.** Salt bridge is defined using a cutoff distance of 4 Å between carboxyl carbon and amine/guanidine nitrogen. From histograms of the distances between the carboxyl carbon and amine/guanidine nitrogen, we note that if acids and bases are involved in salt bridges, then their distance distribution peaks lie within 4 Å.

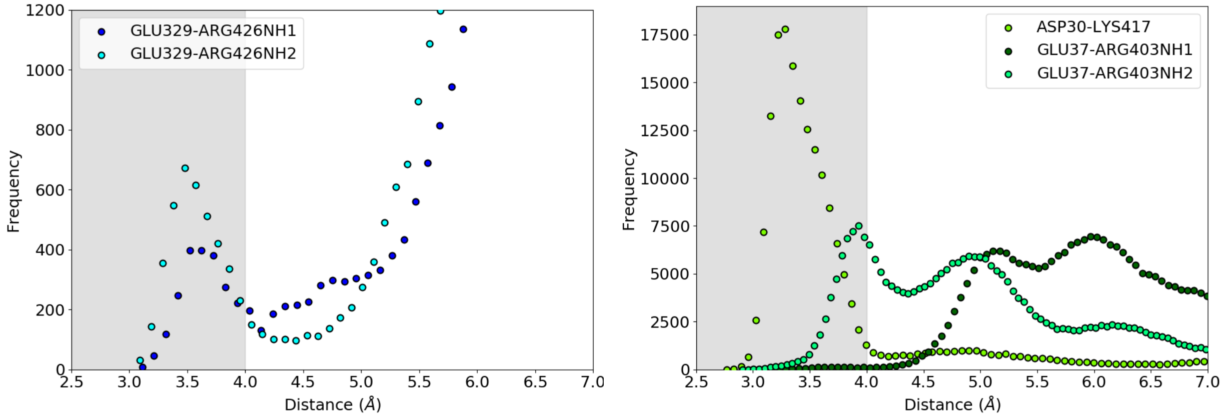

Figure S2: **Comparison of ACE2-spike polar contacts between X-ray structures and MD simulations.** In the MD simulation panel, the colors of the lines connecting the residues indicate their occurrence probability.

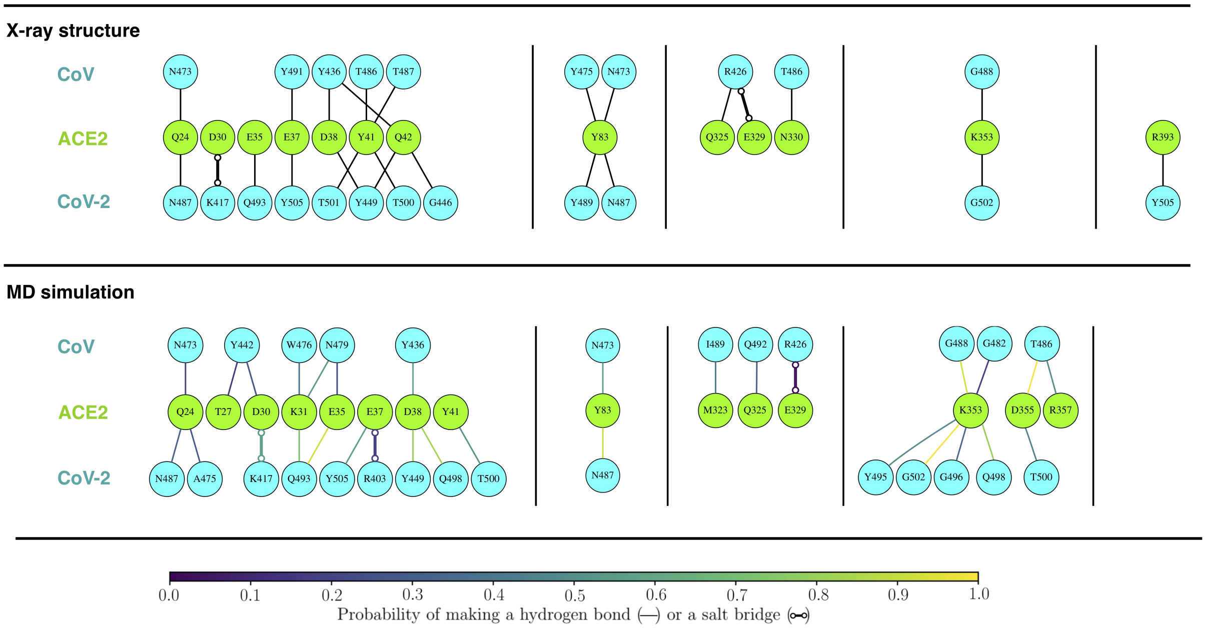

Figure S3: **Hydrophobic contacts between ACE2 and spike proteins of CoV and CoV-2.** The 3D renderings on the left show the surface hydrophobicities of the two spike proteins computed using a normalized consensus hydrophobicity scale.[5] The regions marked by circles are the primary locations on spike that make hydrophobic contacts with ACE2. The plots on the right show time evolutions of hydrophobic contacts. Two hydrophobic residues are considered to make contact if the distances between any of their atoms is less than the sums of their vdW radii.

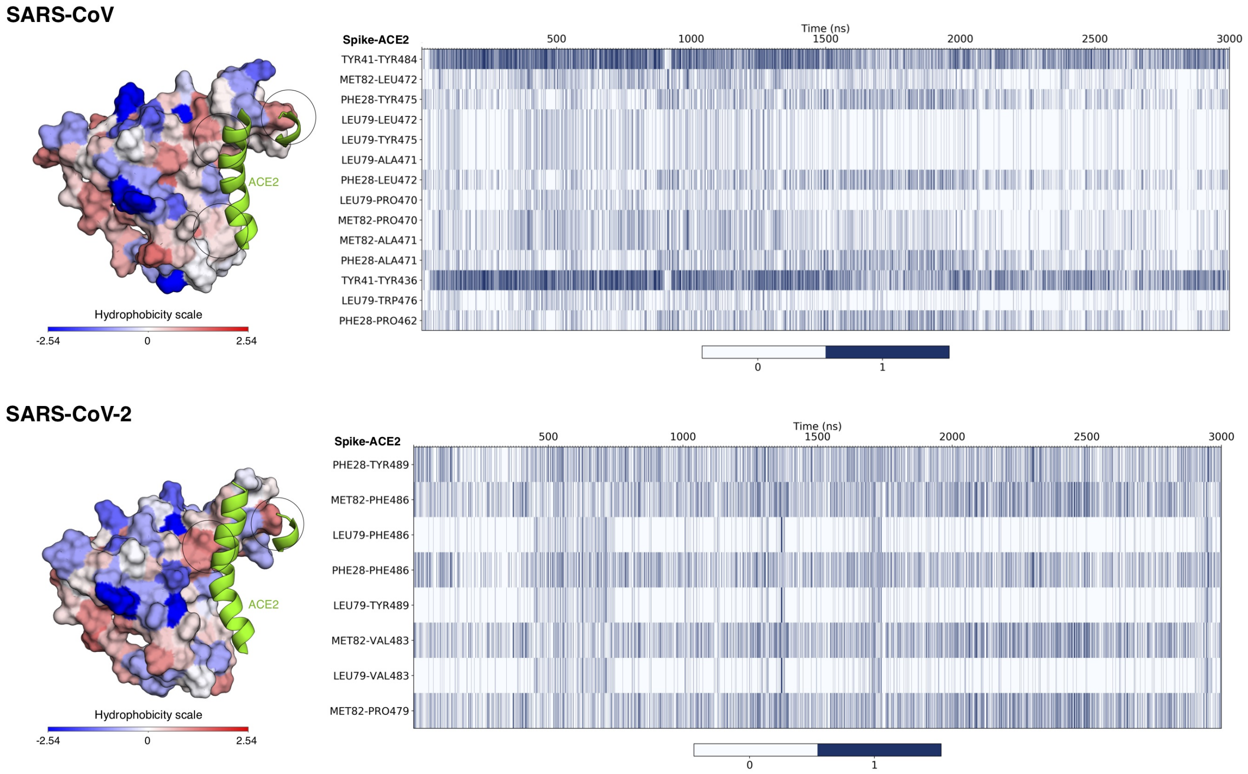

Figure S4: **Effect of CoV spike RBD mutations on its binding free energy with ACE2.** **(a)** Residue binding module of CoV spike protein showing the ten residues at the ACE2-spike interface that are chemically different in CoV-2 spike. Eight of these are mutated to residues present in CoV-2 spike. **(b)** Running averages of $\Delta G$, $\Delta G_{spike}$ and $\Delta G_{spike-ACE2}$ as a function of sampling time. $\Delta G$ is the effect of mutations on spike-ACE2 binding free energy, and is defined as $\Delta G = \Delta G_{spike-ACE2} - \Delta G_{spike}$, where $\Delta G_{spike-ace2}$ is the effect of mutations on the free energy of the spike-ACE2 complex in solution, and $\Delta G_{spike}$ is the effect of mutations on the free energy of isolated spike in solution. **(c)** Running averages of all the $\partial H/\partial\lambda|_{\lambda_i}$ as functions of sampling time.

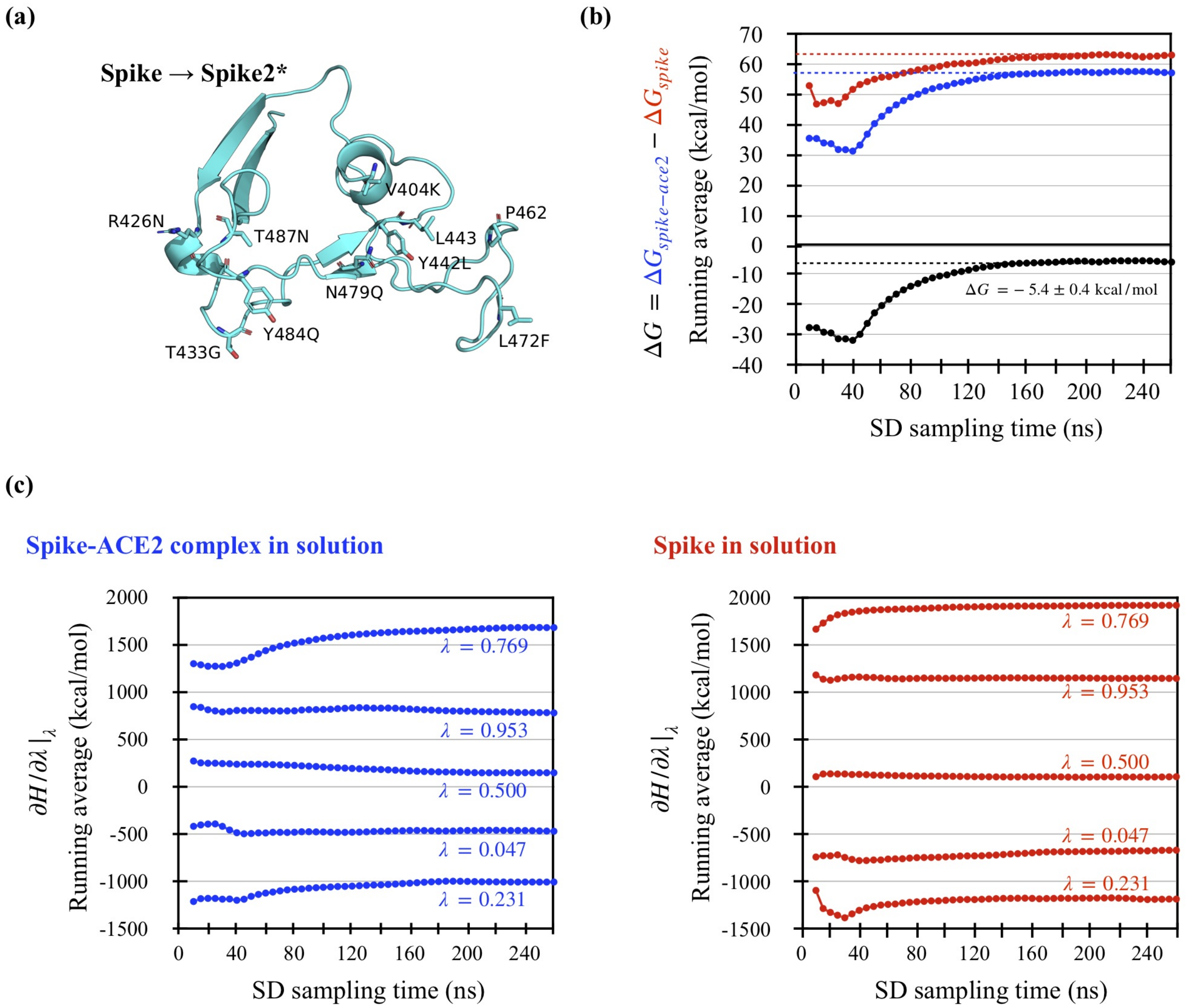
